# randtip, a generalized framework to expand incomplete phylogenies using non-molecular phylogenetic information

**DOI:** 10.1101/2022.01.04.474924

**Authors:** Ignacio Ramos-Gutiérrez, Herlander Lima, Rafael Molina-Venegas

## Abstract

1. The increasing availability of molecular information has lifted our understanding of species evolutionary relationships to unprecedent levels. However, current estimates of the world’s biodiversity suggest that about a fifth of all extant species are yet to be described, and we still lack molecular information for many of the known species. Hence, evolutionary biologists will have to tackle phylogenetic uncertainty for a long time to come.
2. This prospect has urged the development of software to expand phylogenies based on non-molecular phylogenetic information, and while the available tools provide some valuable features, major drawbacks persist and some of the proposed solutions are hardly generalizable to any group of organisms.
3. Here, we present a completely generalized and flexible framework to expand incomplete molecular phylogenies. The framework is implemented in the R package ‘randtip’, a toolkit of functions that was designed to randomly bind phylogenetically uncertain taxa in backbone phylogenies through a fully customizable and automatic procedure that uses taxonomic ranks as a major source of phylogenetic information.
4. Although randtip is capable of automatically generating fully operative phylogenies for any group of organisms using just a list of species and a backbone tree, we stress that the ‘blind’ expansion of phylogenies (using randtip or any other available software) often leads to suboptimal solutions. Thus, we discuss a variety of circumstances that may require customizing simulation parameters beyond default settings to optimally expand the trees, including a detailed step-by-step workflow.
5. Phylogenetic uncertainty should be tackled with caution, assessing potential pitfalls and opportunities to optimize parameter space prior to launch any simulation. Used judiciously, our framework will help evolutionary biologists to efficiently expand incomplete molecular phylogenies and thereby account for phylogenetic uncertainty in quantitative analyses.

## 1. Introduction

The past two decades have seen an explosive interest in incorporating evolutionary history into ecological analyses (Webb et al., 2002; Cavender-Bares et al., 2009; Mouquet et al., 2012), boosting several disciplines such as community ecology (Davies, 2021), macroecology (Lamsdell & Congreve, 2021) and conservation biology (Molina-Venegas et al., 2020). This eco-phylogenetic revolution was driven by the increased availability of molecular information (Sayers et al., 2020) and sophisticated tools for inferring phylogenetic trees (Smith & Walker, 2018), which have lifted our understanding of species evolutionary relationships to unprecedent levels. However, and despite the phylogeny of certain groups is nearly completed (e.g. mammals; Upham et al., 2019), phylogenetic relationships remain vastly uncertain –particularly shallow ones (i.e. infra-family)– for many groups. For example, one of the largest global phylogenies of angiosperm plants published to date includes only ∼12.5% of the species in the group (Janssens et al., 2020), and recent accounts of terrestrial arthropod biodiversity showed that up to 80% of insect species are yet to be discovered (Stork, 2018). These bleak figures suggest that evolutionary biologists will have to tackle phylogenetic uncertainty for a long time to come.

Conscious of the limited extent of molecular phylogenetic information, Rangel et al. (2015) developed a theoretical foundation to systematically account for phylogenetic uncertainty in quantitative analyses. Roughly, the procedure starts with the identification of *phylogenetically uncertain taxa* (PUTs), this is, taxonomic units (e.g. species, subspecies) that are well delineated in the continuum of biodiversity but remain missing from available molecular phylogenies. Then, all acceptable taxonomic, morphological, or behavioral information on the PUTs is used to conservatively define their *most derived consensus clades* (MDCCs), this is, the less inclusive phylogenetic nodes that most certainly contain them. Finally, each PUT is assigned to a random point along one branch of its corresponding MDCC, and the procedure is replicated a high number of times to obtain a distribution of possible trees that can be used in downstream phylogenetic analyses iteratively. While the ‘true’ phylogenetic hypothesis will most certainly remain unsampled, the workflow allows exploring the parameter space, thereby quantifying the extent to which phylogenetic uncertainty has a significant impact in the analyses (e.g. Calatayud et al., 2019; Molina-Venegas et al., 2021). Rangel et al. (2015) accompanied their framework with the software SUNPLIN, a set of algorithms for randomly expanding molecular phylogenies using the aforementioned procedure (Martins et al., 2013).

Although Rangel et al. (2015) suggested that the identification of MDCCs should be based on expert taxonomic evaluation, such knowledge is in practice beyond the reach of most researchers, particularly when dealing with very large phylogenies that often encompass a wide spectrum of taxonomic groups and thousands of species. In a valuable attempt to automatize the identification of MDCCs, Jin & Qian (2019) developed V.PhyloMaker, a R package that generates very large phylogenies of vascular plants. V.PhyloMaker builds on the idea of the classical software Phylomatic (Webb & Donoghue, 2005), which uses a taxonomically informed backbone mega-tree to automatically define MDCCs (genus or family nodes in case the former are not available) and bind the PUTs to the selected clades. Beyond covering features that were already implemented in Phylomatic, V.PhyloMaker provides an option to insert PUTs in randomly chosen nodes below the crown node of the corresponding MDCCs, so that a distribution of possible phylogenies can be generated with relatively little effort (Jin & Qian, 2019)

However, we note that current available tools for the insertion of PUTs, while valuable, have some important drawbacks. For example, V.PhyloMaker uses a pure node-based approach to insert PUTs, and thus the simulations often lead to the formation of polytomies even if a fully bifurcated backbone tree is used. In contrast, SUNPLIN allows the insertion of PUTs along randomly selected branches, but the user must manually set all the MDCCs for the simulations (Martins et al., 2013). V.PhyloMaker circumvents this limitation at the cost of requiring an ‘annotated’ backbone mega-tree (a linkage between all the species represented in the backbone tree and their taxonomic genus and family) that is provided by the developers of the software, and thus the user is forced to use the backbone tree for which the software was implemented –currently the GBOTB.extended tree for vascular plants (see Jin & Qian, 2019)–. Also, the definition of MDCCs on the basis of fixed taxonomic ranks (e.g. genera and otherwise families) might be excessively conservative and hence suboptimal under certain circumstances. For example, large taxonomic families often include taxonomic ranks between the family and genus level that may represent putative MDCCs (e.g. subfamilies, tribes and subtribes in the Asteraceae, Poaceae and Fabaceae plant families). Finally, there are shortcomings that are transversal to all available software for PUT binding, including the disregard of natural paraphyletic groups (Hörandl & Stuessy, 2010) and the impossibility to fully customize the space of branch lengths for the insertion of PUTs among other issues.

Here, we present a completely generalized and flexible framework to expand incomplete phylogenies. The framework is implemented in the R package ‘randtip’, a toolkit of functions that was designed to randomly bind PUTs in backbone phylogenies through a fully customizable and automatic procedure that uses taxonomic ranks as a major source of phylogenetic information. Although randtip is capable of automatically generating fully operative phylogenies for any group of organisms using just a list of species and a backbone tree, we discuss a variety of circumstances that may require customizing simulation parameters beyond default settings to optimally expand the trees, including a detailed step-by-step workflow.

## 2. General workflow

In this section, we describe the general workflow of randtip to expand phylogenies. Roughly, given a list of taxa (typically Linnean binomials) for which a phylogeny is to be obtained and a backbone tree (provided by the user), the software identifies putative MDCCs for the PUTs in the list. MDCCs are defined based on taxonomic ranks, including genus, subtribe, tribe, subfamily, family, superfamily, order and class, and by default the software will select the less inclusive among the available. Once each PUT is assigned to a MDCC, randtip will automatically bind the former to the backbone tree according to the parameters that are set for the simulations, and a phylogeny including all the species in the user’s list is returned (Fig. 1). The workflow can be customized using a variety of parameters that are either passed through the whole simulation or adjusted independently for each PUT (see Appendix for detailed step-bystep examples).

**Figure 1.**
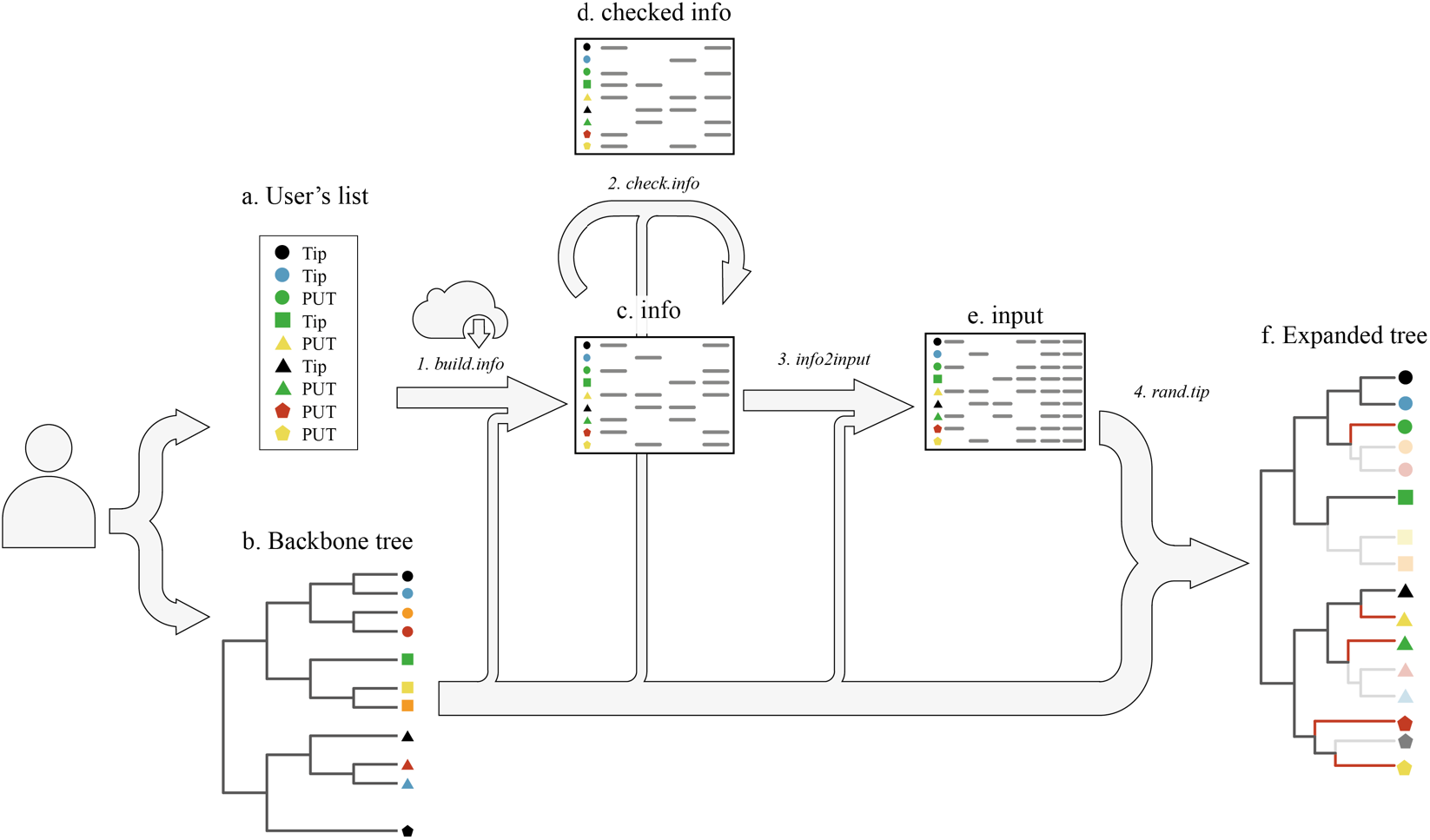
Workflow of the randtip R package. The user provides a backbone phylogeny and a list of species that are to be bound to the former (some of these are already placed in the tree while others represent *phylogenetically uncertain taxa* or PUTs). The function *build*.*info* creates the template *info* and retrieves taxonomic information for the listed species (and for those represented in the phylogeny if the ‘backbone’ mode of randtip is set) from web repositories. The resultant data frame (*info*) can be evaluated with the function *check*.*info*. Once the user has edited *info* according to the particularities of each PUT, the data frame is passed through *info2input* to create the input object for the *rand*.*tip* function, which in turn will expand the backbone phylogeny.

### 2.1. Input files

The workflow of randtip is guided by a data frame R object (hereafter ‘*info*’) and the instructions that are passed through the main function of the package (*rand*.*tip*). The data frame *info* is a template with 21 columns –20 variables of type character or logical plus one integer variable for internal use– that must contain, as a minimum, all the taxa in the user’s list (column 1) and their genus rank (column 2). Optionally, the user may provide supra-generic taxonomic ranks and set parameter values specifically for individual PUTs. For simplicity, we will consider the most common scenario in the ecological literature where the operative taxa represent Linnean binomials (genus and species with or without subspecific epithets), although genus-level phylogenies and taxon lists are also supported. The *info* template can be created automatically using the auxiliar function *build*.*info*, which is fed with species names in a character vector or single column data frame. Besides, *build*.*info* can interact with a suite of taxonomic repositories –currently implemented for ‘ncbi’ (default), ‘itis’, ‘gbif’ and ‘bold’ via the *classification* function of ‘taxize’ R package (Chamberlain et al., 2020)– to retrieve taxonomic information that will be used to identify putative supra-generic MDCCs for the PUTs (note that information to define genus-level MDCCs is intrinsically contained in the scientific names of the species). This can be done by setting the argument ‘find.ranks’ of *build*.*info* to TRUE (default). We recommend providing at least one supra-generic rank (e.g. taxonomic family) for all the species in *info*, which will be used to define MDCCs whenever the genera of the PUTs are missing in the phylogeny (otherwise the PUTs will not be bound). Often the user will need to further edit *info* once the template is automatically created with *build*.*info* (for example, to customize binding parameters for certain PUTs or to amend taxonomic mistakes in web repositories). This can be done directly in R using the auxiliar function *edit*.*info* or exporting the data frame as a spreadsheet (e.g. csv or xlsx) and importing it back into R once all the edits are completed.

The user must provide a backbone phylogeny as a *phylo* R object. Although randtip can identify MDDCs on the sole basis of taxonomic information of the species that are included in both the user’s list and the backbone tree (hereafter ‘taxon list’ mode), MDDCs can also be identified based on taxonomic ranks of all the species that are represented in the latter regardless of their presence in the former (hereafter ‘backbone’ mode). Both approaches have pros and cons, and they will behave identically whenever the genera of the PUTs are minimally represented in the backbone tree. To use the ‘backbone’ mode of randtip, the argument ‘mode’ of *build*.*info* must be set to “backbone” (default) to additionally include all the species in the phylogeny as rows in the *info* data frame, so that their taxonomic information can also be retrieved (if the argument ‘find.ranks’ of *build*.*info* is set to TRUE).

Once the data frame *info* is assembled, we strongly recommend the user to check the incidence of PUTs in the user’s list and their putative MDCCs. This can be done with the auxiliar function *check*.*info*, which will inform on the PUT status of the species, the presence of spelling errors, putative MDCCs, and the phyletic nature of the set of species that are included in each MDCC and share taxonomic ranks (e.g. congenerics, contribals, confamiliars) with the corresponding PUT –hereafter *phylogenetically placed and co-ranked* (PPCR) species–. Also, the tip labels of the backbone tree are checked out for duplicates (e.g. *Ziziphora taurica taurica* and *Ziziphora taurica*) and the ultrametric nature of the tree is assessed. The auxiliar functions *get*.*clade* and *plot*.*clade* can in turn be used to extract and plot respectively any subtree representing putative MDCCs, so that the user can visually explore them using the R graphic window. Exploring MDCCs is particularly recommended to optimize PUT binding, particularly when PPCR species form polyphyletic groups. Alternatively, subtrees can be exported in Newick format to visualize them using auxiliar software such as Dendroscope (Huson & Scornavacca, 2012), which may be very convenient for very large clades. Once the MDCCs are defined and the user has optionally customized parameter values for individual PUTs, the wrapping function *info2input* is fed with the data frame *info* and the backbone tree to create a final dataset that will be passed through the *rand*.*tip* function to expand the phylogeny. This final dataset ensures consistent structure for use in *rand*.*tip* and allows generating as many trees as desired without the need to search for putative MDCCs in *info* repeatedly (this is done by *info2input* just once), which is a computationally intense task.

### 2.2. Selecting MDCCs and binding PUTs

The binding of PUTs is conducted with the function *rand*.*tip*, which includes a variety of parameters that are passed through the whole simulation (Table S1). However, all the parameter arguments of *rand*.*tip* can be adjusted independently for each PUT by editing in the corresponding slots of *info*, which makes the framework completely flexible and customizable.

Randtip will always try to find the less inclusive MDCC for each PUT according to the taxonomic ranks that are provided in *info*, starting from genus level and up to class level until a MDCC for the PUT is found. Regardless of the mode of randtip that is set by the user (‘taxon list’ or ‘backbone’), the software will always first attempt to define genus-level MDCCs as the *most recent common ancestor* (MRCA) of all the species in the backbone tree that are congeneric to the PUTs. However, the definition of MDCCs above the genus level may differ between the two modes of randtip. On ‘taxon list’ mode, supra-generic MDCCs are defined as the MRCA of all the species in the user’s list that are PPCR with the target PUT (e.g. contribals, consubfamiliars, confamiliars). In contrast, the ‘backbone’ mode (default) defines supra-generic MDCCs as the MRCA of all the species in the backbone phylogeny (regardless of their presence in the user’s list) that are PPCR with the target PUT (see Fig. 3 and section 3 for an extended discussion).

By default, *rand*.*tip* will bind each PUT to a randomly selected branch below the crown node of the corresponding MDCC, the probability of being added along any branch being directly proportional to the length of the branch –argument ‘prob’ set to TRUE (default)–. Alternatively, branches can be selected on the basis of equal probability, and in either case the user can decide to add the stem branch of the clade to the pool of candidate branches –if the argument ‘use.stem’ is set to TRUE (default is FALSE)–. The exact point to insert the PUT in the selected branch is sampled from a uniform distribution. Importantly, the extent to which the default behavior of *rand*.*tip* to insert PUTs represents an optimal scenario may depend on the phyletic nature of their PPCR species. These can represent monophyletic (whenever the MDCC is exclusively shaped by species that are PPCR with the target PUT), singleton (terminal branch), paraphyletic (whenever the species that map within the MDCC but are not PPCR with the PUT form either a monophyletic or a singleton group), or polyphyletic (set of PPCR species that do not fit any of the previous categories) groups (see section 4.1 and Fig. 2). For example, the PPCR species of a given PUT could form a polyphyletic clade simply because one of them maps clearly away from the main (monophyletic) cluster of PPCR species –e.g. because the outlying PPCR species is labelled in error (Pentinsaari et al., 2020)–, in which case the default behavior of *rand*.*tip* to bind the PUT (i.e. any branch below the crown node of the largest monophyletic cluster) would be reasonable. In contrast, the polyphyletic nature of PPCR species could be due to ‘intruder’ taxa that map within an otherwise monophyletic cluster, in which case the default behavior of *rand*.*tip* could be suboptimal because the evidence that the largest monophyletic cluster of the group include the PUT may be less clear (Fig. 2). As we discuss in section 4, randtip allows the user to optimize the binding of PUTs according to the specifics of each case.

**Figure 2.**
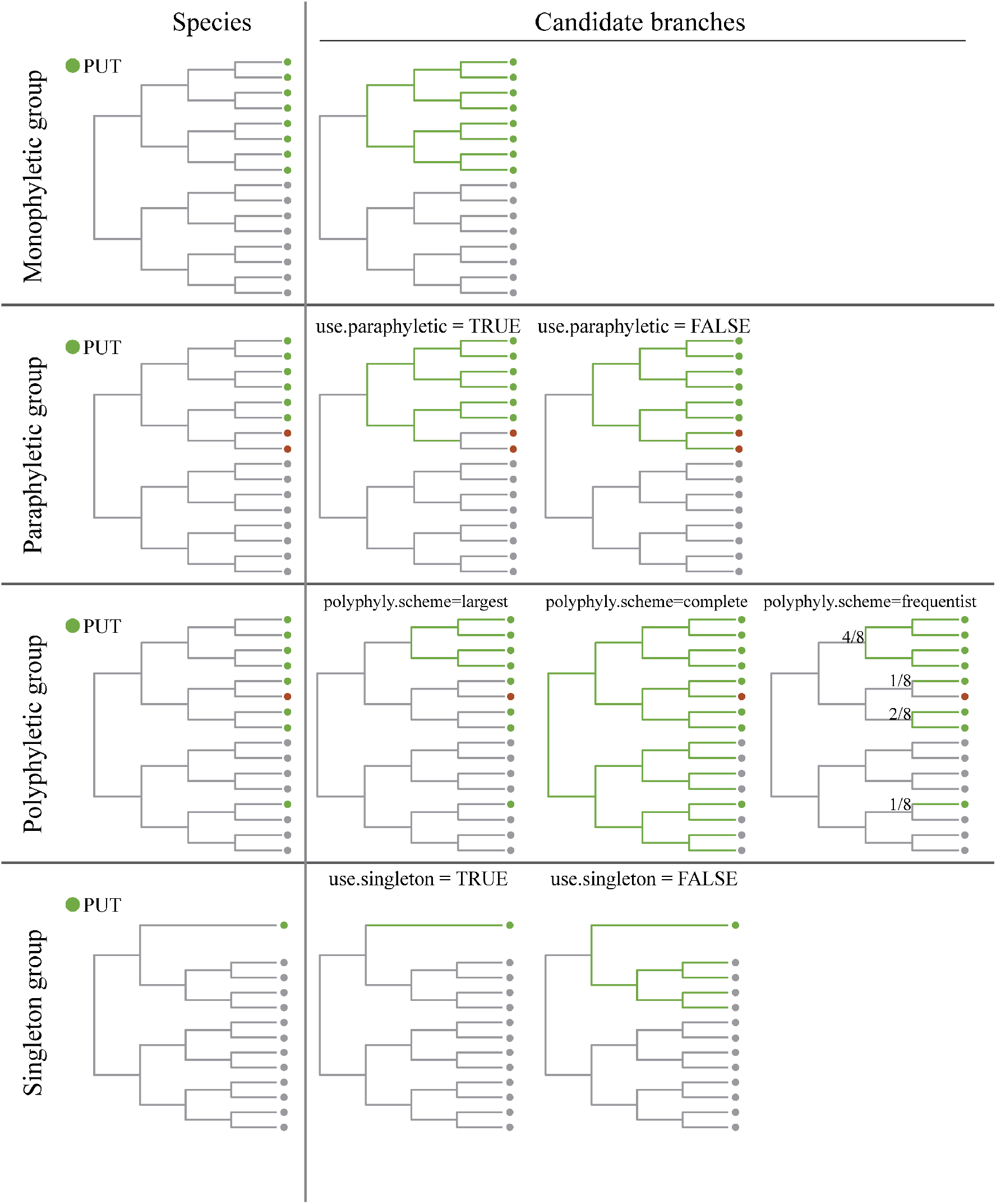
Types of phyletic groups formed by *phylogenetically placed and co-ranked* (PPCR) species (green circle symbols) and possible scenarios for PUT binding within each type. Non-PPCR species are colored in grey, and non-PPCR species that cause an otherwise monophyletic group of PPCR species to be paraphyletic or polyphyletic are colored in red. The candidate branches to bind the PUT in each scenario are colored in green. The fractions close to the phylogenetic nodes indicate the probability for the candidate clades to be selected under the polyphyletic scenario “frequentist”.

It is important to note that the user can always decide to what extent they want to rely on the retrieved taxonomic ranks for the automatic identification of MDCCs. For example, if the taxonomic affiliation of a PUT to a given genus is controversial, the user may edit the data frame *info* to change the genus-rank of the PUT into ‘NA’, in which case randtip will use the taxonomic rank immediately above to find a new MDCC.

## 3. Comparison between ‘taxon list’ and ‘backbone’ modes of randtip

The first decision the user will have to tackle is choosing between the ‘taxon list’ and ‘backbone’ modes of randtip. As we stated earlier, both approaches will perform identically as long as the genera of the PUTs are minimally represented in the backbone phylogeny, yet the definition of supra-generic MDCCs may differ between the two approaches. For example, it might happen that some of the PPCR species of a given PUT (let’s say confamiliars) are missing in the user’s list but are represented in the backbone phylogeny. Thus, in case these PPCR species were phylogenetically external to the confamiliars that are included in the user’s list, the ‘backbone’ mode of randtip would define an older MDCC than ‘taxon list’ (Fig. 3). It follows that the extent of the divergence in the functioning between both modes (whenever a supra-generic MDCC is to be defined) depends on the phylogenetic placement of the PPCR species that are included in the user’s list. In sum, the ‘backbone’ mode works based on the ‘true’ suprageneric MDCCs (but note that these may neither represent the actual MDCCs as the backbone phylogenies are often not fully comprehensive) with the trade-off that it is a more time-consuming approach than ‘taxon list’. In contrast, the latter might define younger supra-generic MDCCs (meaning more restricted parameter space to bind PUTs) under some circumstances (Fig. 3). We recommend considering the ‘backbone’ mode as a first option (default) and use ‘taxon list’ only when there is a low incidence of PUTs requiring supra-generic binding and/or low mismatch in the nodes defining supra-generic MDCCs between both approaches (see Fig. 3 and Appendix for an extended discussion).

**Figure 3.**
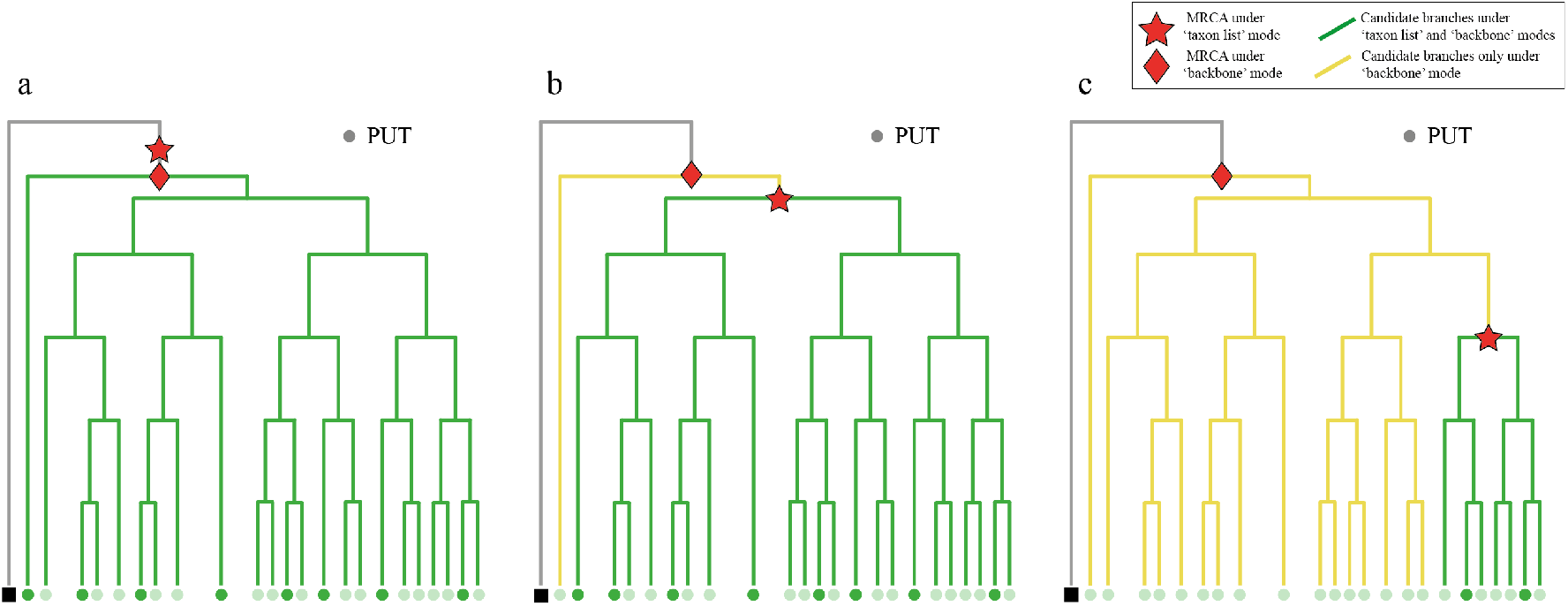
Scenarios of increasing divergence in the performance between the ‘taxon list’ and ‘backbone’ modes of randtip. The circle symbols on the phylogenetic tips represent *phylogenetically placed and co-ranked* (PPCR) species (e.g. confamiliars) of the PUT that is to be bound in each case, and the highlighted ones are those included in the user’s list. The diamond red symbol (hereafter ‘diamond node’) indicates the crown node of the *most derived consensus clade* (MDCC) that is identified for the PUT when taxonomic information is available for all the species in the backbone phylogeny (i.e. under ‘backbone’ mode), and the star red symbol (hereafter ‘star node’) indicates the crown node of the MDCC that is identified when taxonomic information is available only for the species in the backbone phylogeny that are also included in the user’s list (i.e. under ‘taxon list’ mode). In the first scenario (a), the diamond and star nodes are coincident, and thus both modes of randtip will use the same space of branch lengths (in green) to bind the PUT. In the second scenario (b), *the most recent common ancestor* (MRCA) of the subset of PPCR species that are represented in the user’s list includes all PPCR species but one, and therefore the branch subtending the latter (in yellow) will never be selected under ‘taxon list’ mode. In the third scenario (c), a higher number of PPCR species are missing from the user’s list, resulting in a smaller space of branch lengths to bind the PUT under ‘taxon list’ mode. Note that under ‘backbone’ mode, both the green and yellow branches would be candidates to bind the PUT.

## 4. Newly designed features for PUT binding

As discussed above, *rand*.*tip* will by default bind PUTs to randomly selected branches below the crown node of the corresponding MDCCs. However, this default behavior can be modified using a variety of arguments that are implemented in *rand*.*tip*. For example, if the user is not interested in generating a distribution of possible phylogenies but one single tree without randomizing the PUTs, the argument ‘rand.type’ of *rand*.*tip* can be set to “polytomy” (default is “random”) for the function to insert the PUTs as polytomies at the crown nodes of their corresponding MDCCs instead. This is the only binding option that was implemented in the seminal software Phylomatic (Webb & Donoghue, 2005), and it might still be convenient for extremely resource-consuming phylogenetic analyses where using a distribution of possible trees could be computationally prohibitive. Alternatively, the user may want to bind the PUTs following the default behavior of *rand*.*tip* but still inserting *some* of them as polytomies in their corresponding MDCCs. To do so, the user can set the corresponding slots of the column ‘rand.type’ of *info* to “polytomy” (default is NA) while keeping the argument ‘rand.type’ of *rand*.*tip* to “random”.

### 4.1 Polyphyletic, paraphyletic and singleton groups of PPCR species

While PUT randomizations within monophyletic groups of PPCR species will always follow the same scheme (i.e. by default, randomly selected branches below the crown node of the corresponding MDCCs), the user must choose between different scenarios for polyphyletic, paraphyletic and singleton groups. In case MDCCs are shaped by polyphyletic groups of PPCR species, the user must choose between three different scenarios using the ‘polyphyly.scheme’ argument of *rand*.*tip*. If the default option “largest” is set, *rand*.*tip* will pick the largest monophyletic cluster of PPCR species among the available to insert the PUT (less conservative scenario; Fig. 2). If the option “frequentist” is set, *rand*.*tip* will first pick one of the constituent clusters of PPCR species that conform the polyphyletic group, the probability of being selected being proportional to the size of the cluster, and then the PUT will be inserted in the selected cluster. If the option “complete” is set, *rand*.*tip* will bind the PUT to a randomly selected branch below the crown node of the MDCC (most conservative scenario).

In case MDCCs are defined by paraphyletic groups of PPCR species, two different scenarios are eligible. If the argument ‘use.paraphyletic’ is set to TRUE (default), *rand*.*tip* will only bind PUTs to branches so that the paraphyletic nature of the group remains unchanged (Fig. 2). Otherwise, the randomization will be conducted as if the MDCCs were defined by a monophyletic group of species. Importantly, certain taxonomic groups such as the Olacaceae s.l. plant family are paraphyletic (Chase et al., 2016), and thus randomizing PUTs at any point below the crown node of this family (i.e. setting ‘use.paraphyletic’ to FALSE) may result in an excessively conservative parameter space that would encompass almost the entire Santalales order (Malécot & Nickent, 2008).

In case the MDCC of a PUT is defined by one single PPCR species (Fig. 2), *rand*.*tip* will by default bind the PUT to the terminal branch subtending the only PPCR species, and whenever the MDCC is no longer singleton (because at least one PUT was already bound), *rand*.*tip* will consider the entire newly formed clade (same height as the original singleton clade) to sample for candidate branches. We will refer to this procedure as ‘bind-to-singleton’ hereafter. However, if the argument ‘use.singleton’ is set to FALSE (default is TRUE), the parent node of the singleton PPCR species will be defined as the MDCC of the PUT instead (Fig. 2). Although the latter scheme is more conservative than the former, it may lead to suboptimal solutions under some circumstances. For example, the parameter space to randomize a PUT whose MDCC is shaped by one single species that is the only representative of a subfamily in the phylogeny can be drastically increased in case the subfamily is the sister group to the rest of the family. Note that all these parameters can be specifically set for individual PUTs by filling in the corresponding slots of *info*.

### 4.2. Manual definition of MDCCs

Although randtip was conceived to automatize the definition of MDCCs based on taxonomic ranks, the user can manually define MDCCs for the PUTs. This can be done by filling in the corresponding slots of the columns ‘taxa1’ and ‘taxa2’ of *info*. As long as these slots are not set to ‘NA’ (default), the MDCCs of the PUTs will be defined on the basis of this information instead. For example, if the slots ‘taxa1’ and ‘taxa2’ of a PUT are filled in with different species names, the PUT will be bound to a randomly selected branch below the MRCA of the two given species. If both slots are filled in with the same species name, randtip will follow the bind-to-singleton procedure to insert the PUT as sister to the so defined species, and in case the same genus is provided the PUT will be inserted as sister to the clade defined by the MRCA of all the species in that genus.

### 4.3. Respecting monophyletic and paraphyletic clades

By default, *rand*.*tip* will never bind a PUT to a branch that results in breaking the monophyletic or paraphyletic nature of a group (of any taxonomic rank) unless the arguments ‘respect.mono’ and ‘respect.para’ of the function are set to FALSE (default is TRUE). Thus, while previous software followed either approach (e.g. V.Phylomaker always respects monophyletic genera but SUNPLIN does not), randtip offers the user the possibility to choose between both options, either by setting the general arguments of the *rand*.*tip* function or on a customized basis for individual PUTs by filling in the corresponding slots of *info*.

### 4.4. Clumping PUTs

Some genera may not be represented in the phylogeny, and thus their representative species will likely form a polyphyletic group if they are to be bound randomly below the crown node of the corresponding supra-generic MDCC. However, the user could be certain in that a group of congeneric PUTs whose genus is missing in the phylogeny is monophyletic. Thus, if the argument ‘clump.puts’ is set to TRUE (default), *rand*.*tip* will first bind one of the congeneric PUTs, and then the rest will be bound to the former following the bind-to-singleton procedure. Similarly, it may happen that supra-generic taxonomic groups are not represented in the phylogeny, in which case *rand*.*tip* will clump the PUTs as described above and following the taxonomic hierarchy so that the missing taxonomic groups will form monophyletic clusters once all the PUTs are bound. As any other randomization parameter of randtip, the user may decide the PUTs that will be clumped in this way by setting the ‘clump.puts’ option individually in the corresponding slots of *info*.

Trinomials representing infra-specific taxa (e.g. subspecies) are also supported. If ‘clump.puts’ is set to TRUE, *rand*.*tip* will clump PUTs with infra-specific information according to their specific epithets (i.e. second name in the trinomial). To do so, *rand*.*tip* will first check if any of the trinomial PUTs that share specific epithet are represented in the phylogeny. This search also takes into account the type subspecies of the species, which will be detected in either trinomial (e.g. *Ablepharus chernovi chernovi*) or binomial (e.g. *Ablepharus chernovi*) nomenclature. In case one or more PPCR subspecies are found in the backbone tree for a given species, *rand*.*tip* will define a MDCC for the infraspecific PUTs following the standard procedures described in section 4.1. Finally, if none of the trinomials in the group are found, *rand*.*tip* will first bind any of them to the tree, and then all the others will be bound to the former following the bind-to-singleton procedure.

We note that some available phylogenies use, likely in error, both the binomial and trinomial form of a species to label different tips. For example, the GBOTB.tre megatree (Smith & Brown, 2018) includes *Ziziphora taurica taurica* and *Ziziphora taurica* as two different tips. In this case, *rand*.*tip* will randomly select either tip as the actual type subspecies and ignore the other. Although the *check*.*info* function will warn the user about the existence of possible duplicate taxa in the backbone tree, we strongly recommend the user to visually explore tip labelling before expanding any backbone tree.

### 4.5. Non-ultrametric phylogenies

Previous software for PUT binding were conceived to be used with either ultrametric phylogenies (trees with branch lengths where all tips are equidistant from the root) or phylogenies without branch lengths. However, non-ultrametric trees where branch-length are not proportional to time but character distance are also subject of ecological analyses (e.g. Mishler et al., 2014; Mienna et al., 2020). The *check*.*info* function will warn the user in case the backbone phylogeny is non-ultrametric, and *rand*.*tip* will force non-ultrametric trees to be ultrametric –following the nnls method as implemented in ‘phytools’ R package (Revell, 2012)– if the argument ‘forceultrametric’ is set to TRUE (default is FALSE). It is important to note that forcing phylogenies to be ultrametric in this way should not be taken as a formal statistical approach for inferring an ultrametric tree but a method to be deployed whenever a genuinely ultrametric phylogeny read from file fails due to issues related to numerical precision (Revell, 2012). Thus, we strongly recommend the user to visually explore phylogenetic trees that fail the ultrametricity test of *check*.*info* before assuming the failure is due to numerical precision of computer machinery.

If the backbone tree is non-ultrametric and the ‘forceultrametric’ argument is set to FALSE, *rand*.*tip* will simulate the new branch lengths subtending the PUTs by sampling from a uniform distribution *U* (0, L], where L is the length of the longest branch in the backbone tree. In case a backbone phylogeny without branch lengths is provided, *rand*.*tip* will output a phylogeny without branch lengths as well (i.e. topological information only). Thus, the only condition for *rand*.*tip* to accept a phylogeny is that it is rooted.

### 4.6 Customizing a subset of branches to randomize PUTs

The node-based workflow of randtip should suffice to cover most situations in PUT binding exercises. However, the distribution of possible branches for the simulation might not be drawn via MDCCs under some circumstances. For example, taxa of hybrid origin often appear as the sister species or either parent depending on the set of molecular markers that are used for the inference (Wang et al., 2014), in which case phylogenetic uncertainty may pertain to only two singleton putative MDCCs (assuming that the identity of the parents is known and both are represented in the backbone tree). Using the auxiliar function *custom*.*branch*, the user will be able to customize specific subsets of branches to bind PUTs across any segment of the phylogeny.

## 5. Concluding remarks

Randtip is, to our knowledge, the only framework for PUT binding that is completely flexible and generalized, thus addressing several shortcomings of previous designs and offering new opportunities to optimize parameter space in tree expansion exercises. Although randtip can generate fully operative phylogenies using default settings, we stress that accounting for phylogenetic uncertainty should not be conceived as a ‘black box’ procedure for the immediate generation of phylogenies. Indeed, previous studies have documented inaccuracies in the generation of such rough phylogenies due to the ‘blind’ use of software packages (Gastauer & Meira-Neto, 2013). As described above, there is a variety of circumstances that may require customizing simulation parameters for specific PUTs if we are to avoid suboptimal solutions. Beyond providing newly designed tools to expand phylogenetic trees, the framework presented here will help evolutionary biologists to get the most out of the evolutionary information that can be used to guide tree expansion exercises.

## Author contribution statement

RM-V conceived the ideas with inputs from IRG, IRG developed the code with the help of RM-V and HL, RM-V led the writing. All the authors read and commented on the manuscript.

## Acknowledgements

IR-G was primarily supported by the project CM/JIN/2019-005 granted to R.M.-V. (Regional Government of Madrid, Spain), and by the project CGL2017-86926-P (Ministry of Science and Innovation of Spain).

## Glossary of terms

*Phylogenetically uncertain taxon* (PUT): taxonomic unit (e.g. species, subspecies) that is well delineated in the continuum of biodiversity but remain missing from available molecular phylogenies (Rangel et al. 2015).

*Most derived consensus clade* (MDCC): the less inclusive phylogenetic clade that most certainly contains a PUT (Rangel et al. 2015).

*Most recent common ancestor* (MRCA): the youngest phylogenetic node from which a set of taxa descend.

*Taxonomic rank*: level of a group of organisms in the taxonomic hierarchy (e.g. genus, family, order).

*Phylogenetically placed and co-ranked species* (PPCR): set of species that are represented in the phylogeny and share one or more taxonomic ranks with a given PUT (e.g. congenerics, contribals, confamiliars).

## Notes

### Competing Interest Statement

The authors have declared no competing interest.

### Summary of Updates

An acknowledgement section was included as part of the main text

