## Supplementary Table 1 for "randtip, a generalized framework to expand incomplete phylogenies using non-molecular phylogenetic information" (from another version of this preprint: version 1, 2022-01-04)

**Table S1.** Description of the functions implemented in the 'randtip' R package. Default options for variable arguments are listed in the first place.

| Function | Description | Fixed arguments | Variable arguments | Options | Description of variable arguments |
|---|---|---|---|---|---|
| *build.info* | Creates the *info* template and retrieves taxonomic information from web repositories | • Character vector or single-column data frame | tree | NULL, phylo object | A backbone tree of class phylo must be provided if using randtip in "backbone" mode (default) |
| | | | find.ranks | TRUE, FALSE | Retrieve taxonomic information of the species in *info* that will be used to identify supra-generic MDCCs for the PUTs |
| | | | db | "ncbi", "itis", "gbif", "bold" | Taxonomic database from which information is retrieved |
| | | | mode | "backbone", "list" | Whether or not the species in the backbone tree are to be included in *info* |
| | | | interactive | FALSE, TRUE | Whether or not ambiguous species names will be resolved manually by the user or filled in automatically with 'NA' when retrieving taxonomic information |
| | | | genus | FALSE, TRUE | Whether or not a genus-level backbone tree is to be expanded with PUTs representing genera |
| *check.info* | Informs on the PUT status of the species in *info*, spelling errors, putative MDCCs and the phyletic nature of groups of PPCR species | • Data frame *info*<br>• Backbone tree | sim | numeric (0-1) | Similarity threshold to detect possible misspellings on tip labels (default is 0.8) |
| *edit.info* | Auxiliar function to easily edit the *info* data frame | • Data frame *info*<br>• Character vector with the PUTs to be edited or removed | column | string (colname of *info*), NULL | Column of *info* that is to be edited |
| | | | edit | string or logical, NULL | Any allowed value for the column of *info* that is to be edited |
| | | | remove.rows | FALSE, TRUE | Whether or not to remove the corresponding rows of *info* |
| *edit.tree* | Auxiliar function to easily edit the tip labels of the backbone tree | • Backbone tree<br>• Character vector with the tips to be edited or pruned | edit | character vector, NULL | New labels for the specified phylogenetic tips |
| | | | remove.tips | FALSE, TRUE | Whether or not to prune the corresponding tips of the backbone tree |
| *get.clade* | Extracts any subtree representing putative MDCCs | • Data frame *info*<br>• Backbone tree | clade | string (taxonomic group) | MDCC to extract, defined as the MRCA of all the species in the taxonomic groups displayed in *info* |
| *plot.clade* | Plots the clade extracted with *get.clade* and highlights PPCR species in the | • Output of *get.clade* | ppcr.col | string (color) | Color for tip labels corresponding to the species in the taxonomic group indicated in *get.clade* (default is "green") |

| | | | | | |
|---|---|---|---|---|---|
| | corresponding taxonomic group | | nonppcr.col | string (color) | Color for tip labels corresponding to the species not included in the taxonomic group indicated in *get.clade* (default is “blue”) |
| | | | unknown.col | string (color) | Color for tip labels corresponding to the species whose taxonomic group is unknown or missing (default is “grey”) |
| | | | … | … | More plotting arguments passed to the *plot.phylo* function of ‘ape’ R package |
| *info2input* | Wrapping function to create the input data frame for *rand.tip* | • Data frame *info*<br>• Backbone tree | - | - | - |
| *rand.tip* | Binds PUTs into a backbone tree | • Data frame created by *info2input*<br>• Backbone tree | rand.type | “random” or “r”, “polytomy” or “p” | Defines whether PUTs will be bound to randomly selected branches below the crown node of their selected MDCCs or as polytomies instead |
| | | | polyphyly.scheme | “largest” or “l”, “frequentist” or “f”, “complete” or “c” | Defines the procedure to bind PUTs whenever their PPCR species form polyphyletic groups |
| | | | use.paraphyletic | TRUE, FALSE | If TRUE, branches constituting monophyletic or singleton groups of non-PPCR species within the MDCCs of the PUTs are prohibited |
| | | | use.singleton | TRUE, FALSE | If TRUE, PUTs are bound to terminal branches whenever their PPCR species form singleton clades in the phylogeny. Otherwise, they are bound to any branch below the parent node of the singleton PPCR species |
| | | | use.stem | FALSE, TRUE | Whether or not including the stem branches of the clades selected for PUT binding as candidate branches |
| | | | respect.mono | TRUE, FALSE | Whether or not to respect monophyletic groups when binding PUTs above genus level |
| | | | respect.para | TRUE, FALSE | Whether or not to respect paraphyletic groups when binding PUTs above genus level |
| | | | clump.puts | TRUE, FALSE | Whether or not co-ranked PUTs (e.g. congenerics, confamiliars) should be clumped together in the phylogeny in case their taxonomic group (e.g. genus, family) is missing in the tree. Will also clump conspecific PUTs (subspecies) in case none of them are represented in the phylogeny |
| | | | prob | TRUE, FALSE | Whether or not branch selection probability must be proportional to branch length or equiprobable |

| | | | | | |
|---|---|---|---|---|---|
| | | | prune | TRUE, FALSE | Whether or not the newly expanded tree will include only the species in the user's list |
| | | | forceultrametric | FALSE, TRUE | Whether or not the backbone tree will be forced to be ultrametric following the nnls method as implemented in 'phytools' R package |
| | | | verbose | TRUE, FALSE | Whether or not to print information about the flow of the function |
| *put.tip.col* | Set color pattern to distinguish between phylogenetically placed taxa and PUTs when plotting trees with the *plot.phylo* function of 'ape' R package using the "tip.color" argument | • Expanded tree<br>• Backbone tree | placed.col | string (color) | Color for tip labels representing taxa present in the phylogeny (default is "grey") |
| | | | put.col | string (color) | Color for tip labels representing PUTs (default is "red") |
| *custom.branch* | Binds PUTs across customized segments of the phylogeny | • Backbone tree<br>• Data frame *edges* | rand.type | "random" or "r", "polytomy" or "p" | Defines whether the PUT will be bound to the randomly selected branch or as a polytomy in the parent node of the selected branch |
| | | | forceultrametric | FALSE, TRUE | Whether or not the backbone tree will be forced to be ultrametric following the nnls method as implemented in 'phytools' R package |
| | | | prob | TRUE, FALSE | Whether or not branch selection probability must be proportional to branch length or equiprobable |
| *plot.custom.branch* | Auxiliar function for *custom.branch* to visualize candidate branches | • Backbone tree<br>• Data frame *edges* | PUT | NULL, string | Only required for *edges* data frames including more than one PUT. Specifies which PUT's candidate branches must be plotted. |
| | | | candidate.col | string (color) | Colour for candidate branches (default is "green") |
| | | | forbidden.col | string (color) | Color for non-candidate branches (default is "grey") |
| | | | candidate.lwd | numeric | Width of candidate branches (default is 2) |
| | | | forbidden.lwd | numeric | Width of non-candidate branches (default is 1) |
| | | | … | … | More plotting arguments passed to the *plot.phylo* function of 'ape' R package |
