## Supplementary material for "randtip, a generalized framework to expand incomplete phylogenies using non-molecular phylogenetic information": Tutorial (from another version of this preprint: version 1, 2022-01-04)

### Tutorial for randtip R package

#### Contents

**Introduction** **1**

**Package installation** **2**

**Examples** **2**

Example 1 - The ‘backbone’ mode of randtip . . . . . . . . . . . . . . . . . . . . 2

Example 2 - The ‘taxon list’ mode of randtip . . . . . . . . . . . . . . . . . . . . 15

**Manual definition of candidate branches** **18**

**References** **21**

#### Introduction

This tutorial aims to provide a step-by-step user guide to expand incomplete molecular phylogenies using the randtip R package.

Specifically, the tutorial shows how to expand a hypothetical species-level backbone phylogeny with 56 tips using a list of 54 species. Some of the species in the list are already placed in the backbone tree but others are missing, the latter representing phylogenetically uncertain taxa (PUTs).

We focus on hypothetical rather than real world data to cover a wider array of PUT binding situations. The tutorial will soon be available online, and all the data sets required to complete the tutorial are available as part of the randtip R package.

### Package installation

The randtip R package is available as an open-source software hosted on GitHub at github.com/iramosgutierrez/randtip The package can be installed (along with its dependencies) using the `install_github` function in the *devtools* package (Wickham *et al.* 2021).

```
install.packages("devtools")
library(devtools)

devtools::install_github("iramosgutierrez/randtip")
```

The software will be delivered soon as a formal R package so that it can be installed from CRAN using the utility function `install.packages` or the corresponding pathways of R interfaces (e.g. tab 'packages' and then install' if using *R Studio*).

Once the installation is completed the package can be loaded (along with its dependencies).

```
library(randtip)
```

### Examples

#### Example 1 - The 'backbone' mode of randtip

##### 1. Data loading

The package randtip requires the user to provide a backbone phylogeny (R object of class *phylo*) and a list of taxa (typically species and/or subspecies) for which a phylogenetic hypothesis is to be obtained.

The user must ensure that there are no duplicate taxa in the list, and that taxonomic criteria between the later and the phylogenetic tips have been harmonized.

The phylogeny can be loaded into R using the function `read.tree` of *ape* R package (Paradis & Schliep 2019) and the list of species can be loaded as a single column data frame or as a character vector.

```
sp.list <- read.table("/.../specieslist.txt")
back.tree <- read.tree("/.../backbone.txt")
```

If randtip is already installed in the system, the hypothetical example files can be loaded into the working space.

```
sp.list <- mythology$sp.list
back.tree <- mythology$back.tree
```

##### 2. Building the data frame *info*

Once the backbone phylogeny and the list of species have been imported into the working space, the next step is to build the data frame *info*. The most direct and safest way to assemble *info* is using the `build.info` function.

Here, we will follow the 'backbone' mode of randtip (default) to bind the PUTs to the backbone tree (otherwise the argument 'mode' of `build.info` must be set to "list", see 'Example 2' below). Therefore, *info* will include not only the 54 species in the list, but also the species that are represented in the backbone phylogeny. By default, `build.info` will try to retrieve taxonomic information from the web for all the species in *info* (calling to the 'ncbi' repository as default option). However, this procedure makes no sense for our hypothetical example, and thus the 'find.ranks' argument of `build.info` must be set to FALSE (default is TRUE).

Note that retrieving taxonomic information with `build.info` may take some time for very large data sets. As a guideline, downloading taxonomic information for 10585 genera (~75000 species in total) took approximately 5 hours to complete. This timing depends on the number of genera rather than species.

```
my.info.noranks <- build.info(sp.list, tree = back.tree,
                              find.ranks = FALSE,
                              mode = "backbone")
my.info.noranks # print the data frame info on screen
```

Note that the only taxonomic information in the outputted data frame ('my.info.noranks') is genus-level (second column, which has been automatically filled in with the genus of the species in the list and the backbone phylogeny). Thus, in case the genus of a PUT is missing in the backbone phylogeny, the PUT will not be bound. We recommend providing at least one supra-generic rank (e.g. taxonomic family) for all the species in *info*, which will be used to define supra-generic MDCCs whenever the genus of a PUT is missing in the phylogeny.

For the purpose of completing this tutorial, an alternative *info* data frame with taxonomic information for all the species in the list and the backbone phylogeny can be loaded into the working space.

```
my.info <- mythology$info.backbone
my.info # print the data frame info on screen
```

The new *info* data frame ('my.info') includes the supra-generic taxonomic ranks for all the species in the list and the backbone phylogeny that would have been retrieved by `build.info` from the web (should the species in the example represented real taxa), including class, order and family in all cases and subfamily in some cases.

Note that the species that are represented in the backbone phylogeny but are missing in list show hyphens instead of 'NA' from columns $10^{\text{th}}$ to $20^{\text{th}}$. This is because these columns are intended to customize simulation parameters for the species in the user's list representing PUTs. Thus, while we still do not know which species in the list represent PUTs, we can be certain that the species depicted in 'my.info' that are missing in the user's list (i.e. those showing hyphens from columns $10^{\text{th}}$ to $20^{\text{th}}$) do not represent PUTs.

#### 3. Checking the data frame *info*

Once the *info* data frame is created, it is very convenient to check for possible spelling errors. Otherwise, it may happen that species included in the user's list that are also represented in the phylogeny, but misspelled in either object, are misidentified as PUTs.

Also, we strongly recommend the user to check the phyletic nature of the phylogenetically placed and co-ranked (PPCR) species that define putative most derived consensus clades (MDCCs) for the PUTs, so that informed decisions can be made in accordance with the particularities of each case (see section 4).

Finally, it is important to ensure that the backbone tree is ultrametric (in case the tree read from file is genuinely ultrametric) and does not include duplicate taxa. All these checks can be conducted using the function `check.info`, which will output a new data frame with all the information.

```
my.check <- check.info(my.info, back.tree)
```

Our example data set produces three **warning** messages.

```
## There may be misspelling errors in the species list or the
↪  phylogenetic tips. Please, check the TYPO column in the
↪  outputted data frame.
##
## Tips Yetis_abominabilis_abominabilis and Yetis_abominabilis may
↪  represent the same taxon. Please consider removing one of
↪  them.
##
## The backbone tree is not ultrametric.
```

First, the function warns about the possibility of misspelling errors. As such, the column "Typo.names" of the newly created data frame 'my.check' reveals that the species *Gorgona medusi* (represented in the backbone phylogeny) was erroneously typed in the user's list as *Gorgona medusii* (it may also happen the other way around, this is, that a species is misspelled in the phylogeny but correctly typed in the user's list). It is important that the user corrects misspelling errors in 'my.info', because otherwise the species will remain misidentified as PUTs (as shown in the second column of 'my.check'). This can be done directly in R using the auxiliary function `edit.info` or exporting *info* as a spreadsheet (e.g. csv or xlsx) and importing it back into R once the edits are completed.

Here, we will use the function `edit.info` for this purpose. To do so, the user only needs to indicate the name of the column that is to be edited, the corresponding species (as in the first column) and the new information.

```
my.info <- edit.info(my.info, PUTs = "Gorgona medusii",
                     column = "taxon", edit = "Gorgona medusi")
```

Second, the function informs that two phylogenetic tips (*Yetis abominabilis* and *Yetis abominabilis abominabilis*) may represent the same taxon, and thus the user may consider to pick one of them and disregard the other. Otherwise, randtip will choose randomly for the purpose of binding PUTs. Phylogenetic tips can be easily pruned from the tree using the argument 'remove.tip' of the auxiliary function `edit.tree` (which also serves to edit tip labels, see Table S1 in supplementary material), and the corresponding row of *info* should be removed as well. The latter amend can be conducted using the argument 'remove.rows' of `edit.info`.

```
back.tree <- edit.tree(back.tree,
                       tips="Yetis abominabilis abominabilis",
                       remove.tips=TRUE)
my.info <- edit.info(my.info,
                     PUTs="Yetis abominabilis abominabilis",
                     remove.rows=TRUE)
```

Third, the function informs that the backbone tree is not ultrametric. Because we are certain that the tree read from file is genuinely ultrametric (it is simply detected as non-ultrametric due to numerical precision of computer machinery), we will force the tree to be ultrametric later (see section 6).

#### 4. Customizing simulation parameters

By default, randtip will bind the PUTs to the backbone tree using the parameters that are specified in the arguments of `rand.tip` (see section 5). However, using the same set of parameters to bind all the PUTs may lead to suboptimal solutions in many cases.

For example, consider the PUTs *Draco borealis*, *Draco troglodytes* and *Draco wiverny*, whose genus was identified as a polyphyletic group by the `check.info` function (see data frame 'my.check'). We can take a closer look to the less inclusive clade that includes all the species in the genus *Draco* using the functions `get.clade` followed by `plot.clade`.

```
my.clade <- get.clade(my.info, back.tree, clade = "Draco")
plot.clade(my.clade)
```

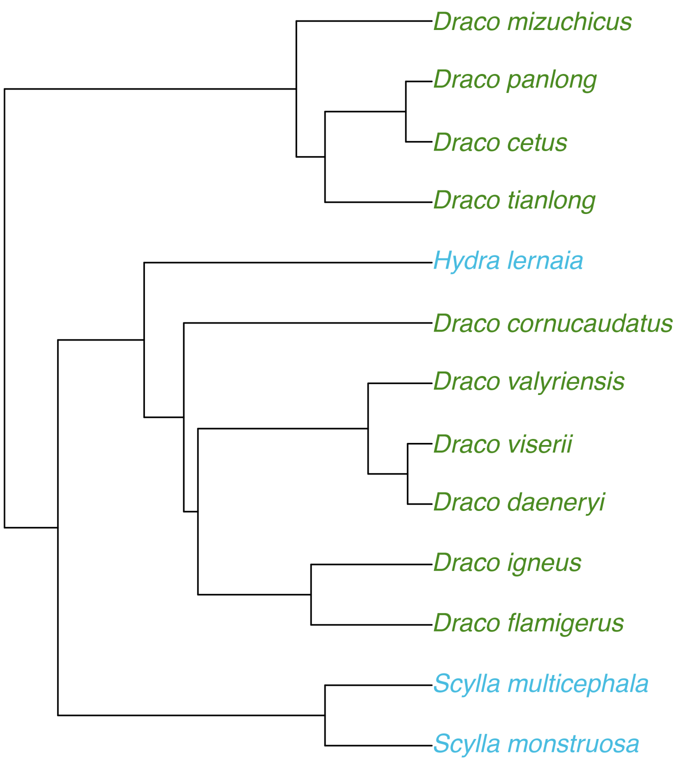


Figure 1: Backbone phylogeny pruned to the less inclusive clade that includes all the species in genus Draco (representative species in green)

Because the congenerics of these PUTs form two monophyletic clusters that are very similar in size (four and six species, respectively; Figure 1), the default behaviour of `rand.tip` for binding *Draco borealis*, *Draco troglodytes* and *Draco wiverny* to this MDCC (largest monophyletic cluster) is risky –the evidence that the largest cluster of Draco most likely include them is weak– and thus a more conservative approach is desirable.

For example, the user may use the “complete” scheme to bind these specific PUTs to a randomly selected branch below the crown node defining the most recent common ancestor (MRCA) of all the species in the genus instead (i.e. root node of the phylogeny displayed in Figure 1). To do so, we can fill in the corresponding slots of *info* (column ‘polyphyly.scheme’) to set the “complete” scheme for these PUTs.

```
DracoPUTs <- c("Draco borealis", "Draco troglodytes",
               "Draco wiverny")

my.info <- edit.info(my.info,
                     PUTs = DracoPUTs,
                     column = "polyphyly.scheme",
                     edit = "complete")
```

It may happen that the user is certain that the MDCC of a PUT does not correspond to any of the taxonomic groups considered by randtip. For example, the MDCC of the PUT *Draco balerion* could be infra-generic (e.g. a taxonomic section within the genus *Draco* including *Draco valyriensis*, *Draco viserii* and *Draco daeneryi*). The user may know that the MRCA of all the species constituting the target taxonomic section in the phylogeny is defined by *Draco valyriensis* and *Draco daeneryi* (the minimum spanning path connecting both species in the tree traverses the MRCA of all the species in the section). Thus, we can fill in the slots “taxon1” and “taxon2” of the corresponding row of *info* with *Draco valyriensis* and *Draco daeneryi* to define an infra-generic MDCC for *Draco balerion*.

```
my.info <- edit.info(my.info,
                     PUTs = "Draco balerion",
                     column = "taxon1",
                     edit = "Draco valyriensis")
my.info <- edit.info(my.info,
                     PUTs = "Draco balerion",
                     column = "taxon2",
                     edit = "Draco daeneryi")
```

Now, consider the PUT Lycanthropus albus, whose genus also forms a polyphyletic group.

```
my.clade <- get.clade(my.info, back.tree, clade = "Lycanthropus")
plot.clade(my.clade)
```

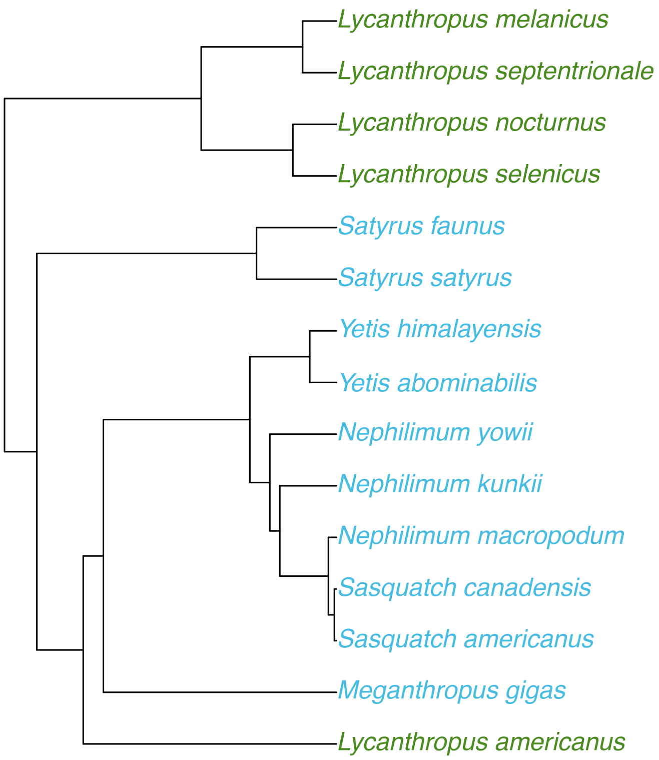


Figure 2: Backbone phylogeny pruned to the less inclusive clade that includes all the species in genus Lycanthropus (representative species in green).

In this case, the polyphyletic nature of the group is due to an outlying species (*Lycanthropus americanus*) that maps clearly away from the main cluster of species in *Lycanthropus* (Figure 2). Therefore, the default scheme "largest" seems adequate (i.e. it is quite likely that the largest cluster of the genus in the phylogeny actually includes *Lycanthropus americanus*) whereas the "complete" scheme could be excessively conservative.

#### 5. Wrapping up

Once we have edited the data in *info* as we see fit (see above), the function `info2input` can be used to create the input object for the `rand.tip` function. This final dataset ensures consistent structure for use in `rand.tip` and allows generating as many expanded phylogenies as desired without the need to search for putative MDCCs in *info* repeatedly, which is a computationally intense task (this is done by the `info2input` just once).

In case `info2input` fails to find a MDCC for a PUT (which will only happen if the genus of the PUT is missing in the backbone tree and no supra-generic taxonomic information is available), the function will return a **warning** message.

```
my.input.noranks <- info2input(my.info.noranks, back.tree)
```

```
## The following taxa were not assigned MDCC and will not be bound
↪  to the tree:
## Grindylowia_yorkii
## Harpia_feminicephala
## Leviathanus_cthulus
## Trolleolus_angmariensis
## Trolleolus_mordoriensis
```

Otherwise, `info2input` will select the less inclusive MDCCs of each PUT.

```
my.input <- info2input(my.info, back.tree)
```

Note that the outputted data frame *my.input* is identical to *my.info* except for the two newly added columns, namely ‘MDCC’ and ‘MDCC.rank’.

‘MDCC’ column shows the taxonomic groups defining the MDCCs to which the PUTs will be bound (‘Tip’ means that the species is already represented in the backbone phylogeny).

‘MDCC.rank’ column depicts the taxonomic rank of the groups.

#### 6. PUT binding

The binding of PUTs in the selected MDDCs is carried out with the `rand.tip` function, which is fed with the output of `info2input` (*my.input*). Most arguments of `rand.tip` are used for defining simulation parameters (and thus they can be customized for individual PUTs via *info*) except for ‘ prune’, ‘forceultrametric’ and ‘verbose’.

By default, `rand.tip` will output a phylogenetic tree including only the species in the user’s list (n = 54 in our hypothetical example) unless ‘prune’ is set to FALSE, in which case the whole expanded backbone phylogeny will be outputted.

In case the tree read from file is detected as non-ultrametric despite being genuinely ultrametric (as in our hypothetical example, see Ramos-Gutiérrez *et al.* (2021) text for an extended discussion on this issue), the user can set the ‘forceultrametric’ argument to TRUE (default is FALSE) to force the tree to be ultrametric.

Lastly, the argument ‘verbose’ allows the user to print the progress of the function on screen (default is TRUE). Here, we will use the function with default settings except for (1) forcing the backbone phylogeny to be ultrametric, and (2) outputting the whole expanded tree (rather than the tree pruned to the species in the user’s list). This will enable us to better visualize the MDCCs that were selected to bind the PUTs (see Figure 3):

```
new.tree <- rand.tip(my.input, back.tree,
                     forceultrametric = TRUE,
                     prune = FALSE)
```

#### 7. Tree visualization

We can visualize the result of the simulation using the `plot.phylo` function of 'ape' R package.

To distinguish between phylogenetically placed species and PUTs, we can set the color pattern of phylogenetic tips using the auxiliary function `put.tip.col` before `plot.phylo`.

Note that visualizing very large phylogenies may require specialized software such as Dendroscope (Huson & Scornavacca 2012).

```
my.tip.col <- put.tip.col(new.tree, back.tree,
                          placed.col = "dark grey",
                          put.col = "red")
plot.phylo(new.tree, tip.color = my.tip.col)
```

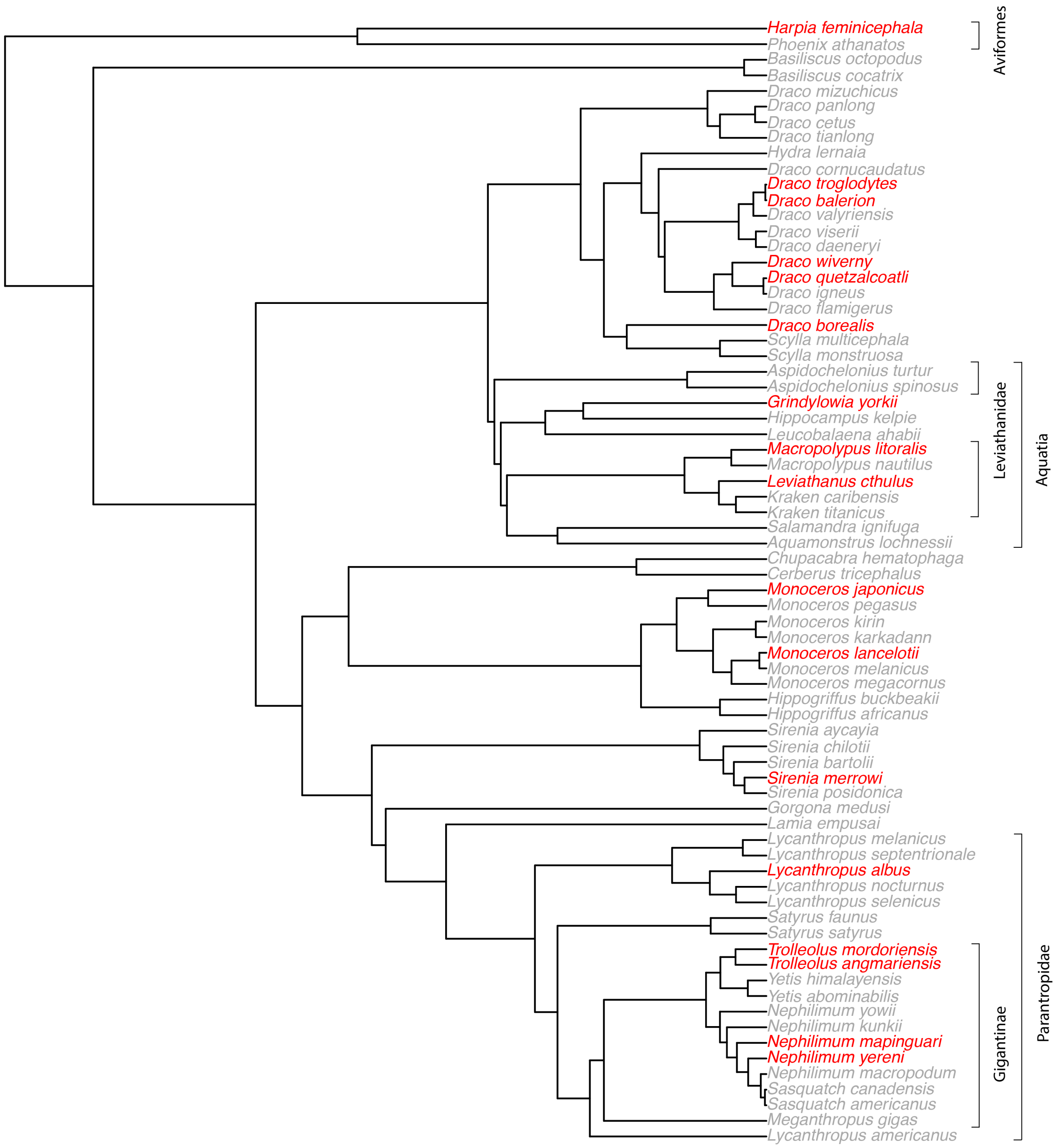


Figure 3: Expanded phylogenetic tree using the 'backbone' mode of randtip. Phylogenetic tips in red represent phylogenetically uncertain taxa (PUTs).

Most of the PUTs were bound to a randomly selected branch below the crown node of their corresponding genus-level MDCCs.

For example, the congenerics of the PUT *Sirenia merrowi* form a monophyletic group, and thus *Sirenia merrowi* is now placed below the crown node of the group (Figure 3).

Scheme 'largest'

In contrast, the congenerics of the PUT *Lycanthropus albus* form a polyphyletic group, and thus it was bound below the crown node of the largest cluster of the genus (default scheme “largest”).

Scheme 'complete'

However, the PUTs *Draco borealis*, *Draco troglodytes* and *Draco wiverny* may not necessarily appear bound below the crown node of the largest cluster of *Draco*, as we specifically set the polyphyletic scheme “complete” to bind these PUTs. Thus, they could have been bound to any branch below the node representing the MRCA of all the species in the genus. For example, in this specific simulation *Draco borealis* was bound as sister to *Scylla*.

Argument use.paraphyletic

The congenerics of the PUTs *Nephilimum yereni* and *Nephilimum mapinguari* form a paraphyletic group, and since the argument ‘use.paraphyletic’ of `rand.tip` was set to TRUE (default), none of the branches subtending the species in *Sasquatch* were considered as part of the parameter space for binding these PUTs. Otherwise, *Nephilimum yereni* and *Nephilimum mapinguari* could have been bound as sister to either species of *Sasquatch*, hence breaking the paraphyletic nature of *Nephilimum*.

Argument respect.mono

In a few cases, the genera of the PUTs were missing in the backbone tree, and thus they were bound to supra-generic MDDCs instead. For example, the MDCC of the PUT *Leviathanus cthulus* was the family Leviathanidae, a polyphyletic group. Note that *Leviathanus cthulus* was bound to the largest cluster of Leviathanidae (default scheme) in such a way that the genera *Macropolypus* and *Kraken* remained monophyletic (Figure 3). This is because the argument ‘respect.mono’ of `rand.tip` was set to TRUE (default).

Argument use.singleton

The less inclusive MDCC of the PUT *Harpia feminicephala* was order Aviformes, which is uniquely represented by the species *Phoenix athanatos* in the backbone phylogeny, and thus the former was bound as sister to the latter because the argument ‘use.singleton’ of `rand.tip` was set to TRUE (default). Otherwise, the parameter space to bind *Harpia feminicephala* would have been substantially larger (any branch where the insertion of the PUT would not compromise the monophyletic or paraphyletic nature of the taxonomic groups represented in the phylogeny).

The PUTs *Trolleolus mordoriensis* and *Trolleolus angmariensis* were bound below the

Argument clump.puts

crown node of the subfamily Gigantinae (less inclusive than Parantropidae), and because the argument 'clump.puts' of randtip was set to TRUE (default), the two PUTs appear clumped together forming a monophyletic group.

Finally, *Grindylowia yorkii* was bound below the crown node of the order Aquatia, its less inclusive MDCC in the backbone phylogeny. Again, note that the monophyletic status of the groups within Aquatia (*Aspidochelonius*, *Macropolypus* and *Kraken*) was kept.

Note that the PUT *Draco balerion* was bound to a branch placed below the MRCA of the species *Draco valyriensis* and *Draco daeneryi*, as we specifically instructed the software to use an infra-generic MDCC to bind this PUT (Figure 3).

#### Example 2 - The 'taxon list' mode of randtip

Now that we are more familiar with the workflow of randtip, we will use the same species list of the previous example to expand the backbone tree using the 'taxon list' mode of randtip.

On 'taxon list mode', randtip defines MDDCs on the sole basis of taxonomic information of the species provided in the user's list, meaning shorter execution times. This is because backbone phylogenies often include thousands of species for which no taxonomic information needs to be retrieved under this mode. However, the definition of supra-generic MDCCs may diverge between both approaches (see Fig. 3 of Ramos-Gutiérrez *et al.* (2021)) which may or may not have an impact on the final tree.

As in the previous example, the first step is building the *info* data frame.

```
my.info.noranks.list <- build.info(sp.list, tree = NULL,
                                   find.ranks = FALSE,
                                   mode = "list")
my.info.noranks.list # print the data frame info on screen
```

The outputted data frame ('my.info.noranks.list') is identical to that generated in the previous example ('my.info.noranks') with the exception that only the species included in the user's list are displayed. Again, we have instructed `build.info` not to retrieve taxonomic information from the web, and thus the only available information is that corresponding to genus rank. For the purpose of completing this tutorial, an alternative *info* data frame including taxonomic information for all the species in the list can be loaded into the working space.

```
my.info.list <- mythology$info.list
my.info.list # print the data frame info on screen
```

Now we can use `check.info`:

```
my.check.list <- check.info(my.info.list, back.tree)
```

Again, we get the same warnings as in the previous example (except for that pertaining to the species that were duplicated in the backbone phylogeny, as we pruned one of them earlier).

Besides, a closer look to the outputted data frame ('my.check.list') reveals that the phyletic status of some groups have changed. For example, the family Leviathanidae is now displayed as monophyletic instead of polyphyletic. This is because the data frame 'my.info.list' only includes taxonomic information for the species in the user's list, and thus any supra-generic taxonomic rank for the species that are represented in the backbone phylogeny but missing in the user's list remains undisclosed. In this case, the two species that conform the small phylogenetic cluster of Leviathanidae (*Aspidochelonius turtur* and *Aspidochelonius spinosus*) are not included in the user's list, which is the reason why the group is now identified as monophyletic (Figure 4).

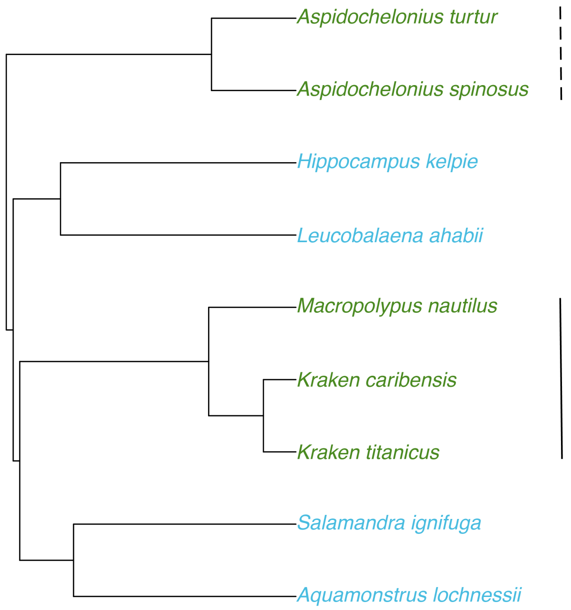


Figure 4: Backbone phylogeny pruned to the less inclusive clade that includes all the species in Leviathanidae (representative species in green). The species next to the solid vertical bar are included in the user's list, whereas those next to the dashed bar are not.

Whether or not the different functioning of the 'backbone' and 'taxon list' modes of randtip for supra-generic MDCCs will have an impact in the expanded tree will depend on the specifics of each situation.

For example, the PUT *Leviathanus cthulus* will always be bound below the crown node of the clade defined by genera Kraken and Macropolypus regardless of the mode of randtip (assuming default settings). This is because under 'backbone' mode, *Leviathanus cthulus* will be bound to the largest cluster of Leviathanidae (n = 3 species; Figure 3), which is the only cluster of Leviathanidae that can be identified under 'taxon list' mode (because the two species in the small cluster of Leviathanidae are not included in the user's list; Figure 4). However, it may have happened that Leviathanidae species not included in the user's list represented the largest cluster of the group in the backbone phylogeny (for example, if the genus *Aspidochelonius* would have been represented by four or more species in the tree), in which case the 'backbone' and 'taxon list' modes would bind *Leviathanus cthulus* to different clades, respectively (if used with default settings).

Finally, it is worth mentioning that both modes of rantip will behave identically whenever the genera of the PUTs are minimally represented in the backbone phylogeny.

#### Manual definition of candidate branches

The clade-based approach of randtip should cover most real-world situations for PUT binding. Yet, the auxiliary function `custom.branch` allows the user to manually define any subset of candidate branches to bind PUTs.

For example, the phylogenetic place for the PUT of hybrid origin *Monoceros x alaricornus* could be any point within the branches subtending the parental species *Monoceros pegasus* and *Monoceros megacornus*, respectively, and such parameter space cannot be specified by one single clade.

To solve this, the user can define the set of candidate branches as an *edges* data frame. The data frame *edges* must contain five columns, each row representing a different set of candidate branches for a given PUT. The first column must include the PUT to which the row refers to. The second and third columns are used to set the older node (MRCA of two given species) and the fourth and fifth ones refer to the younger one. Thus, all the branches traversed by the minimum spanning path connecting the older and younger nodes are selected as candidate branches (the user can add any number of rows as desired).

To define a terminal node (phylogenetic tip) as the younger node, the user must fill in the corresponding slots of the fourth and fifth columns with the corresponding tip label. Inserting the same tip in the four slots will allow binding the PUT as sister to the species represented by the tip. Finally, in case the same pair of species is set for columns 2-3 and 4-5 within a row, the latter will define all branches below the MRCA of the two species as candidates.

In order to ensure that candidate branches have been correctly encoded in *edges*, the user can use the auxiliary function `plot.custom.branch` to visually explore the selected space of branch lengths.

```r
edges <- mythology$edges
plot.custom.branch(new.tree, edges, cex = 0.7,
                   candidate.lwd = 4, forbidden.lwd = 1.5)
```

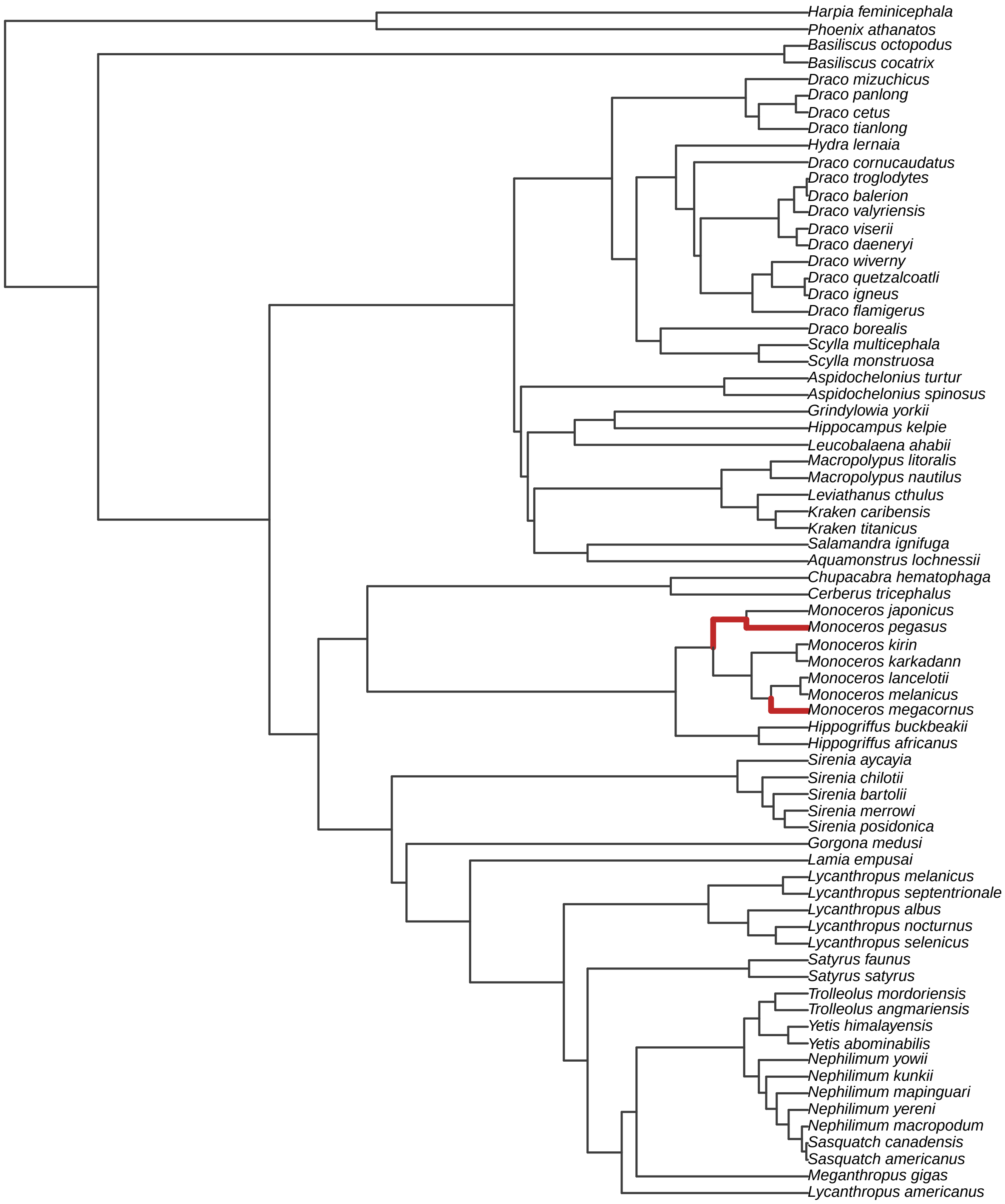


Figure 5: Phylogeny with candidate branches for the insertion of *Monocerus x alaricornus* highlighted in red.

And now we can further expand the phylogeny that was generated earlier.

```
new.tree2 <- custom.branch(new.tree, edges)
plot.phylo(new.tree2, cex = 0.7)
```

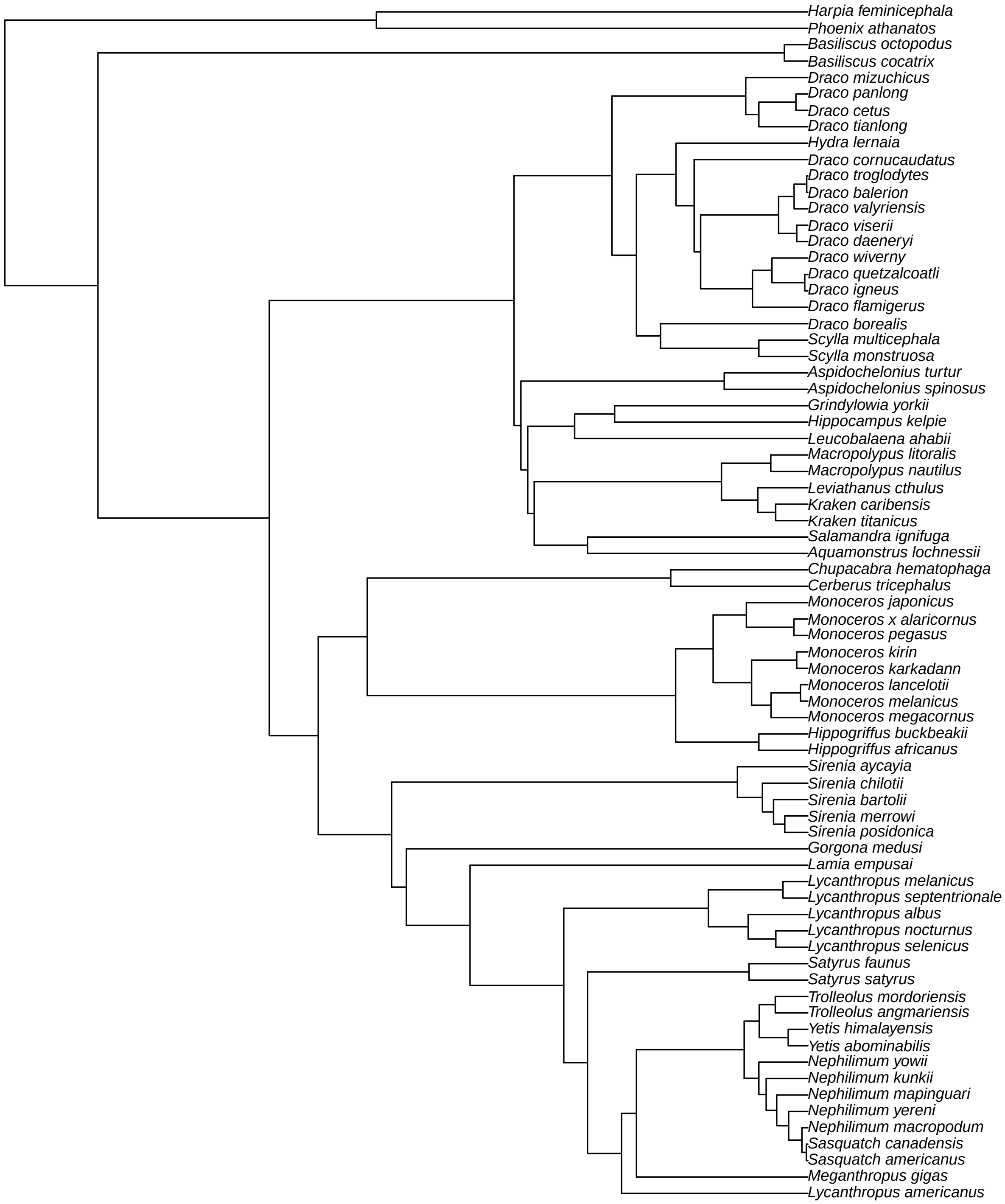


Figure 6: Phylogeny after the insertion of *Monocerus x alaricornus*.
